# The observation of explicit and implicit visuomotor cues can drive predictive motor control

**DOI:** 10.1101/664854

**Authors:** Guy Rens, Marco Davare

## Abstract

Recent studies have highlighted that the observation of hand-object interactions can influence perceptual weight judgements made by an observer. Moreover, observing explicit motor errors during object lifting allows individuals to update their internal sensorimotor representation about object weight. Embodying observed visuomotor cues for the planning of a motor command further enables individuals to accurately scale their fingertip forces when subsequently lifting the same object. However, it is still unknown whether observation of a skilled lift is equally able to mediate predictive motor control in the observer. Here, we tested this hypothesis by asking participants to grasp and lift a manipulandum after observing an actor’s lift. The object weight changed unpredictably (light or heavy) every third to sixth trial performed by the actor. Participants were informed that they would always lift the same weight as the actor and that, based on the experimental condition, they would have to observe skilled or erroneously performed lifts. Our results revealed that the observation of both skilled and erroneously performed lifts allows participants to update their internal sensorimotor object representation, in turn enabling them to predict force scaling accurately. These findings suggest that the observation of explicit as well as implicit visuomotor cues are embodied in the observer’s motor repertoire and can drive changes in predictive motor control.

## Introduction

Skilled hand movements are essential throughout our daily life. It has been well established that dextrous object manipulation not only relies on tactile feedback but also on anticipatory sensorimotor mechanisms. Performing hand-object interactions allows internal object representations to be formed. In turn, these internal sensorimotor representations can be retrieved to enable anticipatory planning of digit forces for future object manipulations (E.g. see Johansson & Westling, 1988). It has been argued that predictive force scaling requires an association between intrinsic object properties, for example size or texture, and the object weight, which are experienced by visual and tactile feedback respectively (Baugh, Kao, Johansson, & Flanagan, 2012). In addition, other research groups have demonstrated that object weight is not only perceived via somatosensory inputs but can also be retrieved through vision and that visual weight judgements are associated to the actual object weight (Bingham, 1987; Runeson & Frykholm, 1981). Finally, it has been established that the object lifting phase conveys critical information for mediating weight judgements: observers mostly rely on the duration of the lifting movement for generating weight perception (Hamilton, Joyce, Flanagan, Frith & Wolpert, 2007; Shim & Carlton, 1997).

The influence of action observation on both weight perception and lift performance was first investigated by Meulenbroek and colleagues: They demonstrated that, when both the actor and subject had an incorrect weight prediction, lifting performance errors made by the subject are reduced, but not eradicated, after observing the actor making typical lift errors (Meulenbroek, Bosga, Hulstijn, & Miedl, 2007). In addition, it was shown in a more recent study by Uçar and Wenderoth that observation of different types of hand movements can alter grip force generation during object grasping: Prior to grasping an object, subjects were asked to observe an actor either touching or squeezing an object. The latter condition led subjects to produce larger grip forces (Uçar & Wenderoth, 2012). Finally, it has been demonstrated that when individuals observe grasping errors, they are able to differentiate object weight based on kinematic cues and, in turn, to scale their fingertip forces more accurately in upcoming trials (Reichelt, Ash, Baugh, Johansson, & Flanagan, 2013). Although these studies have shed light on how action observation can mediate anticipatory motor control in the observer, they only focused on observation of explicit hand-object interactions (different movements [Uçar & Wenderoth, 2012] or salient movement errors [Meulenbroek et al., 2007; Reichelt et al., 2013]) and not on more subtle, implicit, skilled performance of hand movements.

To our knowledge, only a few studies have compared how observing erroneous and skilled object interactions can mediate predictive force scaling. For example, using the size-weight illusion, Buckingham and colleagues highlighted that predictive force scaling in the observer is significantly better after observing erroneous compared to skilled lifting. That is, when participants had to lift a large, but unexpectedly light object for the first time, those who observed typical overestimation errors on the same object would predict the actual weight more accurately (Buckingham, Wong, Tang, Gribble, & Goodale, 2014). Interestingly, when investigating how corticospinal excitability (CSE) was modulated during lift observation, Buckingham et al. found that only during the observation of skilled lifts, CSE was modulated by object size: CSE modulation was significantly higher in response to the observation of a skilled lift of the larger object compared to the smaller one. However, during observation of erroneous lifts on the same objects, the effect of object size on CSE modulation was eradicated (Buckingham et al., 2014). As such, it seems that, when observing skilled object lifting, object size is the critical factor for extracting object weight and driving CSE changes; while when observing erroneous lifts, kinematic cues, not size, have a predominant effect. As a result, it seems plausible that when a lifting error is observed, the unexpected object kinematics drive individuals to shift their attention towards the object kinematics and not size, improving the observer’s predictive force scaling and altering the underlying CSE modulation.

In the current study, we aimed to specifically investigate whether observation of skilled object lifting can drive changes in internal sensorimotor representations when a similar action observation strategy is used for both erroneous and skilled lifts. For this reason, we emphasised on three factors considering the aforementioned studies: (1) we used objects that are identical in appearance to exclude that size and other visual cues could be used to predict object weight. (2) Similarly to the study of Reichelt et al., participants were familiarized to the experimental protocol and object weights. (3) In contrast to the study of Reichelt et al., subjects were informed that they would have to focus on the observation of either skilled or erroneous object lifting. We argue that these factors would allow participants to better understand the task goal and to specifically focus on the actor’s movement kinematics during both action observation conditions. Even though kinematic differences during skilled movements are far more subtle than during erroneously performed movements (e.g. see Buckingham et al., 2014), we hypothesized that observation of skilled lifts can mediate predictive force scaling similarly to observation of explicit lift errors.

## Methods

### Participants

14 subjects (6 males and 8 females; mean age = 19.7 ± 2.9 years) were recruited from the student body of KU Leuven to participate in the current study. All participants were right-handed (self-reported), had normal or corrected-to-normal vision, were free of neurological disorders and had no motor impairments of the right upper limb. The study was conducted in accordance with the declaration of Helsinki and was approved by the local ethical committee of the Faculty of Biomedical Sciences, KU Leuven. Subjects were financially compensated for their participation. Data of one participant were rejected after the data analysis stage due to high inconsistencies in grasping pattern throughout the experiment.

### General procedure

Subject and actor were comfortably seated opposed to each other in front of a table (for the experimental set-up see figure 1A). Participants were required to grasp and lift a manipulandum (see’Data acquisition’) that was placed in front of them (1) either repeatedly (‘SOLO condition’) or (2) in turns with the actor (‘dyadic conditions’). Participants and actor used their entire right upper limb to reach for the manipulandum and were asked to grasp it with the thumb and index finger only (precision grip). Subjects and actor were required to lift the manipulandum smoothly to a height of approximately 3 cm and to keep the grasp-and-lift movement consistent throughout the entire experiment. Additionally, subjects and actor were required to place their hand on a predetermined resting position on their side of the table between trials, at a distance of approximately 25 cm from the manipulandum. This was done to ensure consistent reaching movements across trials. Each trial initiated with a neutral sound cue (‘start cue’) indicating that the movement could be initiated. Trials lasted 4 seconds to ensure that subjects and actor had enough time to reach, grasp and lift the manipulandum smoothly at a natural pace. Inter-trial interval was approximately 5 s during which the weight of the manipulandum could be changed. A transparent switchable screen (Magic Glass), placed in front of the participants’ face, became transparent at trial onset and turned back to opaque at the end of the trial. The screen remained opaque during the inter-trial interval.

**Figure 1.**
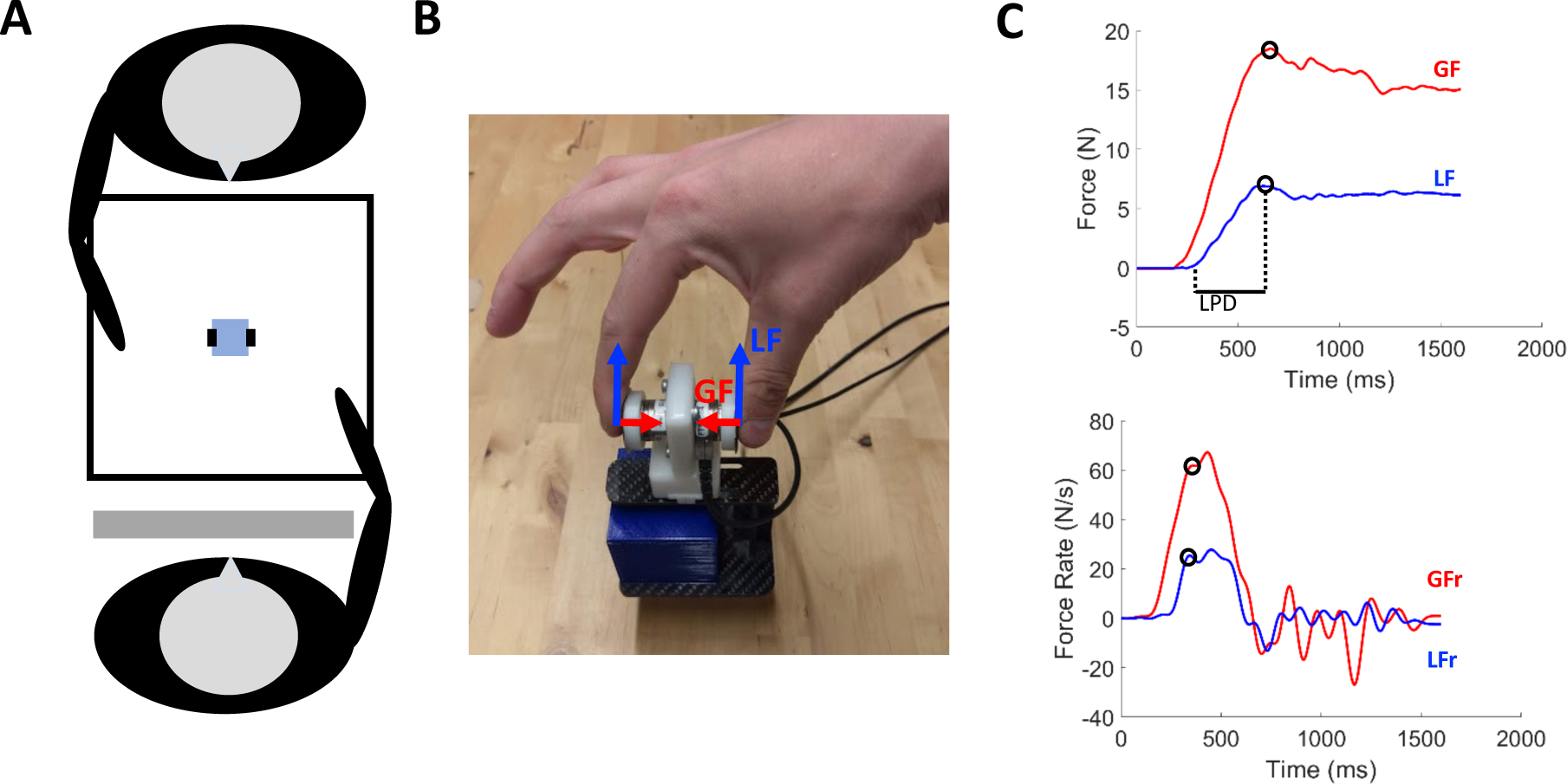
A. Experimental set-up: The participant and actor are seated opposite each other at a table on which the manipulandum was positioned and a screen was placed in front of the participant’s face. **B**. Photo of the grip-lift manipulandum used in the experiment. Load force (LF: blue) and grip force (GF: red) vectors are indicated. **C**. GF and LF typical traces (upper) and their derivatives (lower) for a skilled lift. Circles denote first peak values used as parameters. Loading phase duration (LPD) is indicated on the upper panel.

### Experimental conditions

We used an experimental set-up similar to the study of Reichelt et al. (Reichelt et al., 2013). Participants always performed the solo condition first in order to be familiarized with the experiment. After this condition, subjects performed two dyadic conditions, i.e. erroneous lift observation (‘EO’) and skilled lift observation (‘SO’). Each dyadic condition was performed two times in a counterbalanced order within and across subjects.

#### Solo condition (‘SOLO’)

Participants repeatedly lifted the manipulandum themselves, therefore performing all trials. The weight of the object changed between 1.5 N (light, ‘L’) and 6.2 N (heavy, ‘H’) after a pseudo-random amount of trials of the same weight. The number of trials per weight sequence (i.e. sequential lifts of the same weight) varied randomly between 3 and 6 trials. Thus participants could not predict when the weight change would occur based on the number of lifts. Subjects completed 8 transitions from each weight to the other (i.e. from 1.5 N to 6.2 N and vice versa). This provided 8 trials per weight transition which were used to familiarize participants, assess baseline sensorimotor memory effects (for example see: Johansson & Westling, 1984) and use for comparison with the dyadic conditions.

#### Dyadic conditions

Between the end of the SOLO condition and the start of the first dyadic condition, subjects were instructed on lifting errors i.e. incorrect scaling of fingertip forces due to wrong estimation of object weight. They were told that in the dyadic conditions they would have to lift the manipulandum in alternation with the actor and that the object weight presented in their trial would always be identical to the weight lifted by the actor in the preceding trial. It was also mentioned that the object weight would always change first for the actor and then would be the same for the subject. Finally, subjects were asked to avoid making lifting errors and, importantly, they were told to use cues from the actor’s movement to estimate object weight. However which movement cues could be relevant or which strategy could be used were not discussed. After receiving the task instructions, participants performed the two dyadic conditions. As in the SOLO condition, there were 8 transitions from one weight to the other after a pseudo-random amount of trials. During the dyadic conditions each weight sequence consisted of an even amount of trials between 6 and 12. As such, both actor and participants lifted the manipulandum between 3 and 6 trials within each weight sequence (i.e. the same amount as in the SOLO condition for each person).

Because each dyadic condition took twice the amount of trials in comparison with the SOLO condition, both dyadic conditions were divided into two blocks with a break in between them. This was done to prevent fatigue affecting observation and movement performance. Dyadic block order was counter-balanced within and between subjects. Although both dyadic conditions consisted of two separated blocks, data is presented pooled per condition. In the SO condition, the actor always scaled his fingertip forces correctly to the weight that was presented to him. As a result, the subject could only extract information about object weight by observing skilled lifts. In the EO condition, the actor incorrectly scaled his fingertip forces when the new weight was presented. This lifting error was made only in the first trial after the weight change. In all other trials of the same weight sequence of the EO condition, the actor would perform a skilled lift of the manipulandum. Thus in the EO condition, participants could perceive a weight change by looking for lifting errors. Importantly, the lifting error made by the actor was intentional due to the experimental set-up (see: ‘data acquisition’). Lastly, one of the authors (GR) served as an actor for all experiments.

### Data acquisition

A grip-lift manipulandum consisting of two 3D force-torque sensors (Nano17, ATI Industrial Automation, Apex, NC, USA) was attached to a custom-made carbon fibre basket in which different objects (cubes) could be placed (For an example of the manipulandum see Fig. 1B). The total weight of the manipulandum was 1.2 N. The graspable surface (17 mm diameter and 45 mm apart) of the force sensors was covered with fine sandpaper (P600) to increase friction. The objects were 3D-printed cubes of 5 × 5× 5 cm, filled with different amounts of lead particles to create weights of 0.3 N (‘light’) and 5.1 N (‘heavy’), therefore the total weight were respectively 1.5 N and 6.3 N for the light and heavy weight. To exclude all visual cues about weight, cubes were hidden under the same paper cover. It is noteworthy that cubes were changed manually between each trials (even for trials without weight change) to ensure participants could not use any sound cues to predict weight changes. Second, given the actor was responsible for changing cubes between trials, he always knew what weight would be presented in the upcoming trial. Therefore, the over- and underestimation lift errors related to object weight were made intentionally by the actor and not by a wrong prediction of object weight. Custom-made scripts were compiled in MATLAB (Mathworks) for both data acquisition and processing.

### Data analysis

Force signals were sampled in 3 dimensions at 1000 Hz and smoothed using a 4th order, zero-phase lag, low-pass Butterworth filter with a cut off frequency of 15 Hz. Grip force (GF) was defined as the exerted force (on the force sensors) perpendicular to the normal force. Load force (LF) was defined as the exerted force parallel to the normal force (Fig. 1B). GF and LF were computed as the sum of the respective force components exerted on both sensors. Additionally, grip force rate (GFr) and load force rate (LFr) were calculated by computing the first derivative of GF and LF. Finally, we calculated the loading phase duration (LPD) by measuring the latency between LF onset (LF>0.05 N) and an approximation of object lift off (LF>0.95 * total object weight) (Fig. 1C). Peak force rate values, not peak force values, are presented in the results as it has been demonstrated that these force parameters are a reliable indicator of predictive force scaling (Gordon, Forssberg, Johansson, & Westling, 1991; Johansson & Westling, 1988). These force parameters were compared based on only the first and second trials after the weight change for both subject and actor as it has been demonstrated that individuals adapt to the actual object weight after one trial (Gordon, Westling, Cole & Johansson, 1993). As such, these trials allowed us to investigate (1) the baseline for over- and underestimation of object weight by subjects during the SOLO condition, (2) the movement kinematics of the actor in the EO and SO condition and (3) whether the EO and SO conditions alter the typical over- and underestimation of object weight and could mediate accurate predictive force scaling.

### Statistical Analysis

For statistical analysis of peak force rate values, we normalized the data of all subjects; i.e. the peak values and LPD of each trial were divided by the peak values and LPD of the last trial in the same weight sequence (i.e. sequential lifts of the same weight). For example: If the subject had to grasp 5 heavy weights repeatedly, all parameters of these 5 trials were divided by the parameter value recorded in the fifth trial of the same sequence. The first 4 trials are expressed as a ratio to the fifth trial and the fifth trial would have a value of 1 for each parameter. If any of the measured parameters in the last trial of the weight sequence was an outlier relative to this condition (value larger or smaller than mean ± 2 SD’s) then the entire sequence of weight repetitions was discarded. We chose to compute ratios based on the last trial of a weight sequence because the last trial can be considered as the most skilled due to the repetition of lifts of the same weight (Reichelt et al., 2013). Secondly, some participants altered their general force pattern over time during the experiment although they were informed to maintain a consistent grasping pattern. Using this procedure, the over- and underestimations of object weight are always expressed in relation to the force pattern of skilled lifting during that specific time point and take these potential changes over time into account.

We performed repeated-measures ANOVAs to investigate differences in the weight change trials between conditions. We used 2 within-subject factors: LIFT NUMBER (the first trial after weight change and the second trial after weight change) and CONDITION (SOLO, SO, and EO). Importantly, when investigating the actor’s force parameters, we included the last trial of each weight sequence in the factor LIFT NUMBER in order to investigate the actor’s consistency. ANOVAs were performed separately for heavy-to-light and for light-to-heavy weight changes. Comparisons of interest exhibiting statistically significant differences (p ≤ 0.05) were further analysed using the Holm-Bonferroni test. All data presented in the text are given as mean ± standard error of the mean (SEM).

## Results

We aimed to investigate whether action observation can drive changes in internal sensorimotor representations, which would further translate into changes in predictive motor control. To address this issue, we compared 3 conditions. In the solo condition (‘SOLO’), participants repeatedly lifted the objects for familiarization purposes and to assess baseline sensorimotor memory effects caused by an unexpected weight change. In the dyadic conditions, participants lifted series of objects in alternation with an actor. Subjects were informed that they would always have to lift the same object weight as the actor. For this reason, subjects could use observed kinematics to perceive object weight and consequently update their internal sensorimotor representation. In the error observation condition(‘EO’), the actor would make a typical lifting error when the weight would change from light to heavy (i.e. ‘undershoot’) or from heavy to light (i.e. ‘overshoot’). The actor would then correctly scale his fingertip forces in the following trials. In the skilled lift observation condition (‘SO’), the actor would always apply correct fingertip forces. These two action observation conditions allowed us to investigate whether individuals respond differently to error vs. skilled actions in order to plan their own motor command following an unexpected object weight change.

### Actor’s lifting force parameters

The actor only lifted the objects during action observation trials. In EO, we expected that the first trial after a weight change would differ significantly from the following lifts of the same weight (i.e. explicit lift error). In SO, we expected all trials, including the first lift after a weight change, to be performed with comparable force parameters (i.e. skilled lift).

For all force parameters and both weight changes, except pLFr (*F*_*(1,13)*_ *= 0.54, p = 0.47, Fig. 3A*) and LPD (*F*_*(1,13)*_ *= 2.71, p = 0.12*, Fig.3C) for the heavy-after-light weight changes, both main effects of CONDITION and LIFT NUMBER as well as the CONDITION X LIFT NUMBER interaction were significant *(all F-values>8.39, all p-values < 0.01, Figures 2-3)*.

**Figure 2.**
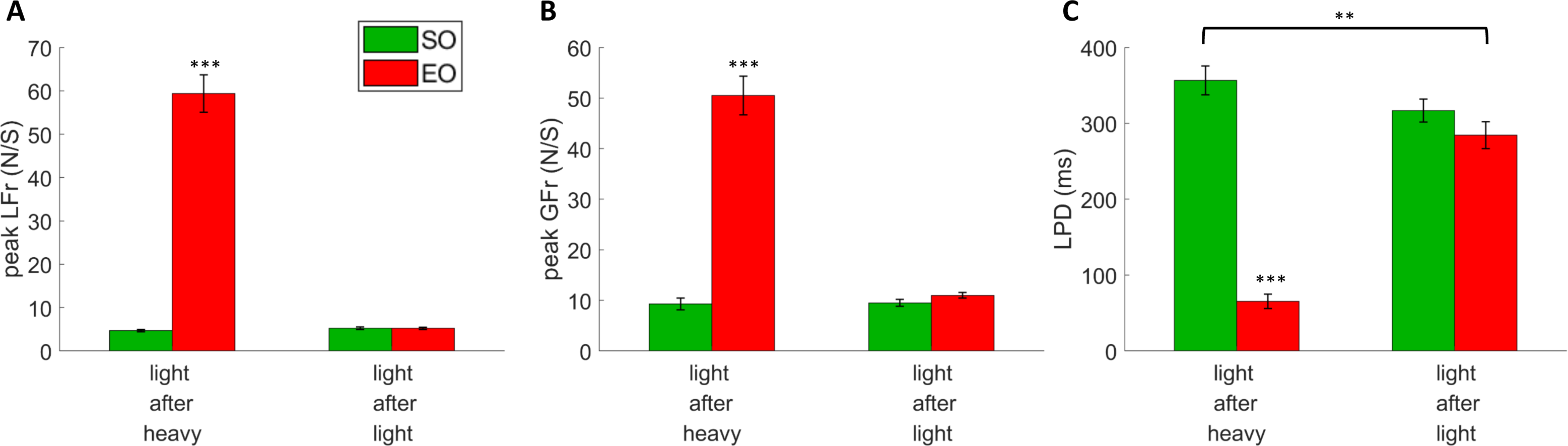
Averaged lift performance for the actor pooled across all participants for light object lifts either after a heavy lift (left bars) or a light one (right bars). A, B and C show averaged data for peak LF rate (LFr), peak GF rate (GFr) and LPD, respectively. Green bars represent lifts performed by the actor in the skilled condition and red bars represent the error condition. All data is presented as the pooled mean ± SEM. ***p>0.001, **p>0.01, *p>0.05. When the asterisk is placed above one bar only, this indicates that this condition significantly differed from all others.

**Figure 3.**
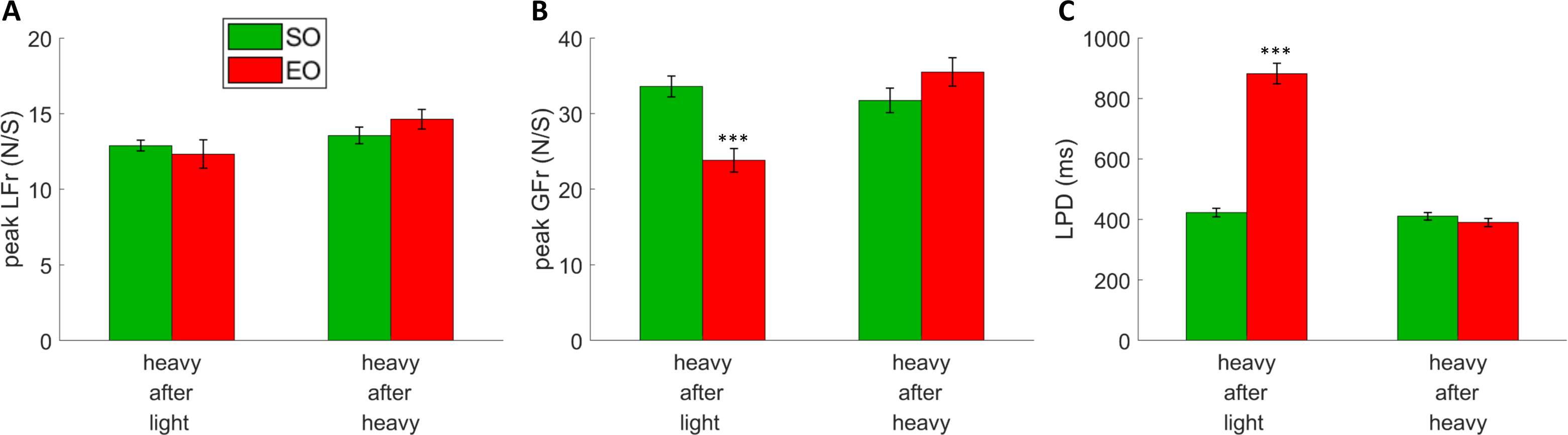
Averaged lift performance for the actor pooled across all participants for heavy object lifts either after a light lift (left bars) or a heavy one (right bars). A, B and C show averaged data for peak LF rate (LFr), peak GF rate (GFr) and LPD, respectively. Green bars represent lifts performed by the actor in the skilled condition and red bars represent the error condition. All data is presented as the pooled mean ± SEM. ***p>0.001, **p>0.01, *p>0.05. When the asterisk is placed above one bar only, this indicates that this condition significantly differed from all others.

Table 1 represents the actor’s lifting performance, pooled across all participants. The force pattern used by the actor in the first trial after the weight change in the EO condition was significantly different from all other trials of both conditions: Post-hoc analyses for all force parameters revealed that, except for pLFr in the heavy-after-light condition, that the first trial after the weight change in the EO condition differed significantly from all other trials of both conditions (Figures 2 and 3).

**Table 1.** Mean peak force rates and loading phase duration for the lifting movements performed by the actor. The actor’s lifting performance, pooled across participants, is presented as mean ± SEM. Values represent the peak load force rate (pLFr; N/s^2^), peak grip force rate (pGFr; in N/s^2^) and loading phase duration (LPD; in ms) for the first, second and last trials of the weight sequence blocks of both dyadic conditions (SO: skilled lift observation, EO: erroneous lift observation). As the actor only made lifting errors during the first trial of the EO condition, this trial should significantly differ from the parameters of all other trials of both the SO and EO condition. *indicates whether the first trial of the EO condition differs significantly from the same parameters of all other trials.

|  |  | Skilled lift observation |  |  | Erroneous lift observation |  |  |
| --- | --- | --- | --- | --- | --- | --- | --- |
|  |  | Trial 1 | Trial 2 | Last trial | Trial 1 | Trial 2 | Last trial |
| **Light**<br>**after**<br>**heavy** | pLFr | 4.70 ± 0.26 | 5.27 ± 0.33 | 5.03 ± 0.29 | 59.38 ± 4.48* | 5.22 ± 0.25 | 5.19 ± 0.31 |
|  | pGFr | 9.30 ± 1.19 | 9.52 ± 0.74 | 8.59 ± 0.52 | 50.50 ± 3.97* | 11.01 ± 0.58 | 11.53 ± 1.36 |
|  | LPD | 356.63 ± 19.78 | 316.65 ± 15.75 | 315.10 ± 16.44 | 65.46 ± 10.05* | 284.4 ± 18.5 | 306.16 ± 15.38 |
| **Heavy**<br>**after**<br>**light** | pLFr | 12.89 ± 0.38 | 13.56 ± 0.58 | 12.30 ± 0.46 | 12.32 ± 0.98 | 14.63 ± 0.67 | 12.03 ± 0.5 |
|  | pGFr | 33.59 ± 1.43 | 31.75 ± 1.69 | 30.90 ± 2.22 | 23.82 ± 1.61* | 35.50 ± 1.97 | 31.20 ± 2.56 |
|  | LPD | 422.83 ± 14.57 | 410.39 ± 13.00 | 453.37 ± 15.45 | 881.8 ± 35.34* | 389.95 ± 14.2 | 474.45 ± 16.08 |

In order to explain the lack of effect for pLFr in the heavy-after-light weight changes, we further looked at the time-to-peak of pLFr. Post-hoc analysis of the significant interaction effect (CONDITION X LIFT NUMBER *F-value = 4.83, p-value = 0.02)* revealed that the pLFr time-to-peak value was significantly longer for the first trial after the weight change in the EO condition compared to all trials in all other conditions. These analyses highlight that the actor’s lifting performance was explicitly different for the first trial after a weight change compared to the following trials, thus providing reliable lifting error cues to the observer.

It is noteworthy that the lifting errors were made artificially by the actor as he always had prior knowledge of the object weight. For this reason, errors were exaggerated in comparison with natural lifting errors on similar weight differences (for example see: Reichelt et al., 2013). Hamilton and colleagues showed that strong deviations in loading phase duration influence weight perception in the observer (Hamilton et al., 2007), therefore it is plausible that subjects are still capable of deriving object weight based on these artificial lifting errors. In addition, the EO condition was essentially added to replicate the findings of Reichelt and colleagues (Reichelt et al., 2013). The main purpose of the current study was to investigate whether skilled lift observation can mediate sensorimotor memory. Importantly, our data revealed that the actor was consistent throughout the performance of skilled lifting as there were only two cases for which LPD values were significantly different (light-after-heavy: first SO trial vs. second EO trial, *p < 0.01*; heavy-after-light: second vs. last trial in the EO condition, *p > 0.01)*.

### Observers’ lifting force parameters: light-after-heavy weight changes

The left panel of Figure 4 shows the averaged force profiles of the first trial after the weight changed from heavy to light in a typical subject. When a subject scales his fingertip forces in anticipation of a heavy object, more force than required (overshoot) will be applied to lift the light object adequately (Johansson & Westling, 1988). In addition, it has been demonstrated that after observing a lift error, individuals are able to immediately scale their fingertip forces accurately (Reichelt et al., 2013). Accordingly, Figure 4 reveals that the subject was able to downscale force parameters after observing a lift error compared to the SOLO condition. It is noteworthy that the subject was also able to apply the correct force scaling after observation of a skilled lift (SO). For data analysis purposes, we only included the first and second trials following a light-after-heavy weight change. Considering that we processed the data using ratio values (see ‘Methods’), force scaling overestimation corresponds to peak force rates with ratios larger than 1. As expected for light-after-heavy weight changes, these effects are the opposite for the loading phase duration: A ratio value smaller than 1 indicates a faster increase in force generation thus resulting in a shortened loading phase duration.

**Figure 4.**
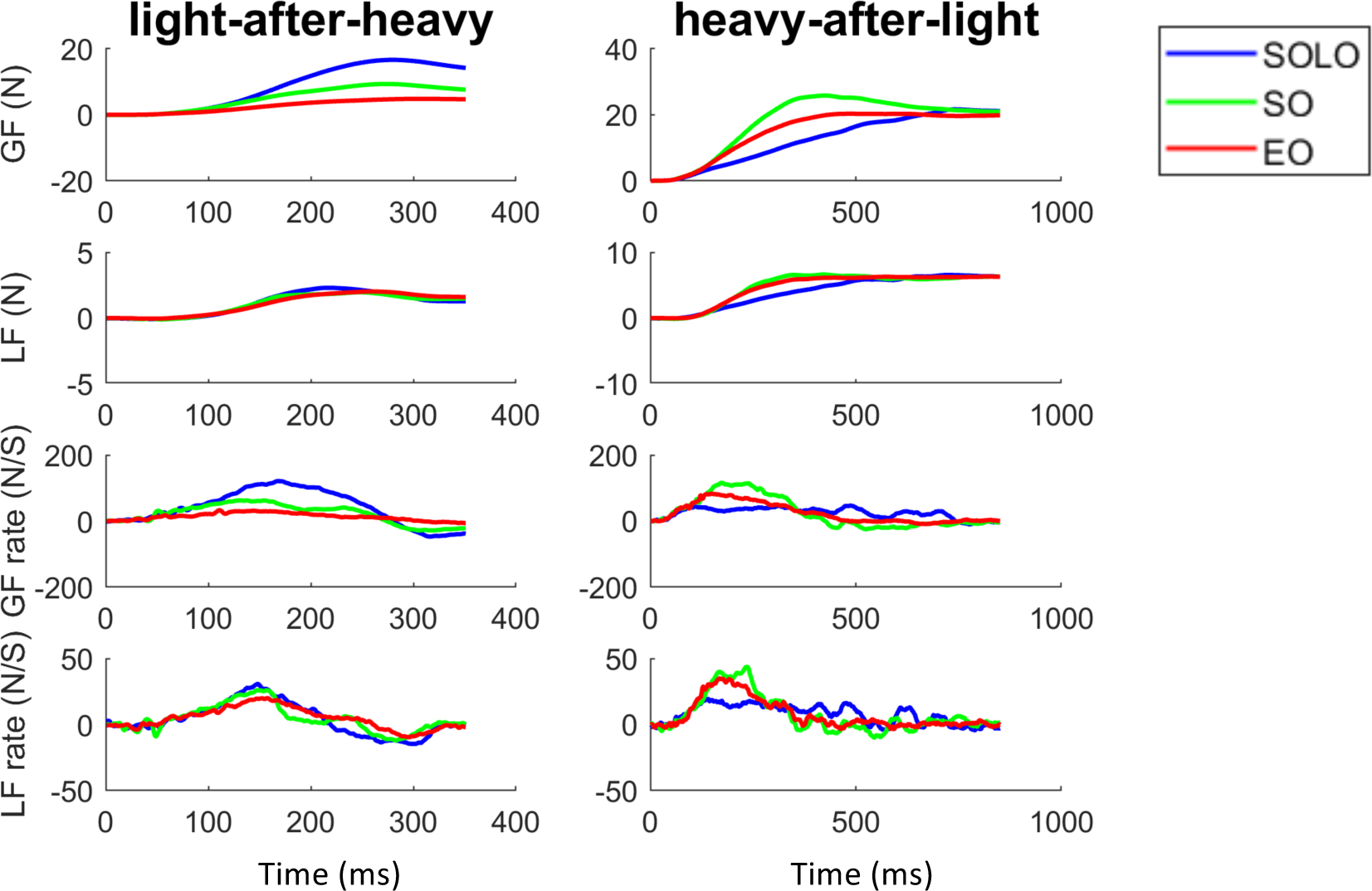
Typical traces showing the evolution of the different force profiles over time for one subject for the three conditions: grasping with incorrect weight expectations (SOLO), grasping after observing a skilled lift (SO) and grasping after observing a lifting error (EO). From top to bottom: Grip force (GF), load force (LF), grip force rate (GFr) and load force rate (LFr) for light-after-heavy (left panel) and heavy-after-light (right panel).

#### Load force rates

Repeated-measures ANOVA for load force rates revealed that both main effects were significant (*both F-values>5.65; both p-values < 0.05)*. The interaction effect was not significant (*F*_(2, 22_ *= 0.19, p = 0.83)*. Firstly, as can be seen in Figure 5A, post-hoc analysis of the significant main effects revealed that participants scaled their load forces with a significantly improved accuracy after observing lifting errors in comparison with the SOLO condition *(p < 0.01)*. However, given that there was no significant difference between the SO and SOLO conditions and between the SO and EO conditions *(p> 0.31)*, this indicates that the SO condition is likely to mediate predictive force scaling as well albeit to a lesser extent than the EO condition. Finally, considering the significant main effect of LIFT NUMBER, it is clear that participants were able to predictively scale fingertip forces with increased accuracy in the second trial after the weight change (*p = 0.01)*.

**Figure 5.**
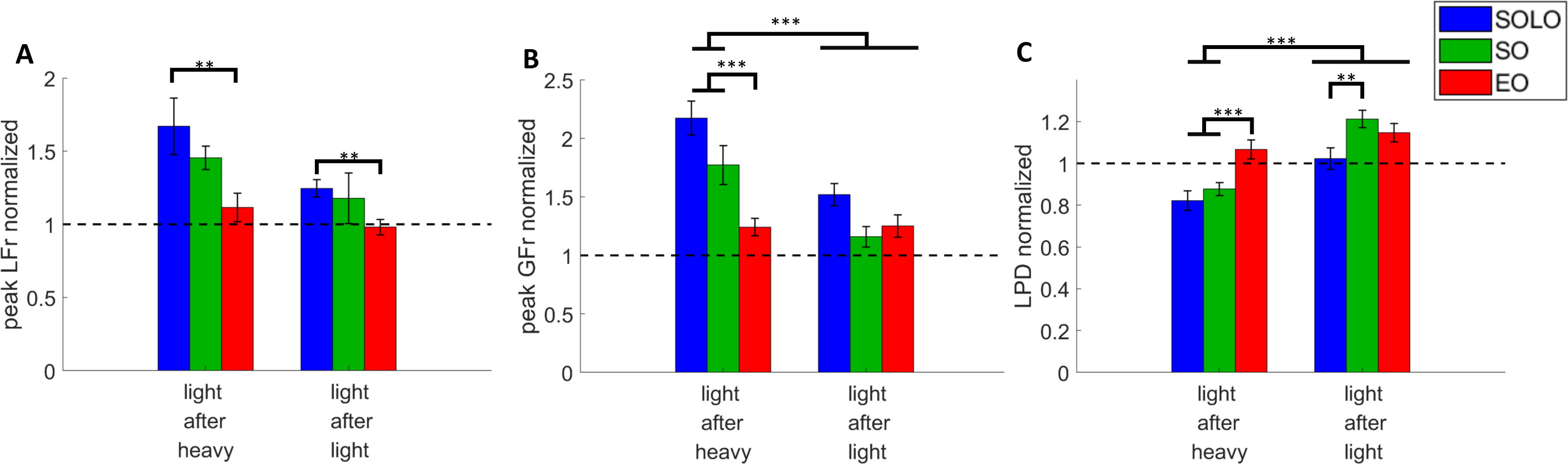
Light-after-Heavy weight changes: Subject group averages for the first and second trials after the weight changed from heavy-to-light for the three conditions [grasping with incorrect weight expectations (SOLO), grasping after observing a skilled lift (SO) and grasping after observing a lifting error (EO)]. **A**. Peak load force rates, **B**. Peak grip force rates and **C**. Loading phase durations. All data is represented as a ratio (normalized to skilled lifting, light-after-heavy and light-after-light divided by the last light-after-light lift of the same weight sequence block). A ratio>1 for peak grip force rates and peak load force rates (and a ratio > 1 for loading phase durations) indicates that subjects overestimated object weight. All data is presented as the pooled mean ± SEM. *** p > 0.001, ** p > 0.01, * p > 0.05.

#### Grip Force rates

The analyses for peak grip force rate revealed that both main effects and the interaction effect were significant (*all F-values>7.34; all p-values < 0.01)*. As can be seen in Figure 5B, it is noticeable that participants scaled their grip forces with the highest accuracy after observing errors (Ratio = 1.16 ± 0.08) in comparison with both the SO (Ratio = 1.77 ± 0.77) and the SOLO (Ratio = 2.17 ± 0.14) conditions (*both p-values < 0.001)*. In addition, the difference between the SO and SOLO conditions neared significance *(p = 0.11)*, indicating, in line with the findings for LFr, that the observation of skilled lifting might be able to mediate predictive force scaling. Finally, for all conditions, participants were increasingly accurate in the second trial after the weight change as the analysis revealed no significant differences between the SOLO, SO and EO second trials *(All p-values = 1)*.

#### Loading phase duration

Repeated-measures ANOVAs for the loading phase duration revealed that both main effects and the interaction effect were significant *(all F-values>8.18, all p-values < 0.05)*. In Figure 5C, it is noticeable that participants had a significantly shorter loading phase duration in the SO (Ratio for first trial of SO = 0.88 ± 0.03) and SOLO condition (Ratio for first trial of SOLO = 0.82 ± 0.05) in comparison with all other trials of all conditions (*all p-values < 0.05*). This indicates that the observation of lift errors allowed participants to lift the object accurately (Ratio for first trial of EO = 1.07 ± 0.05). Finally, participants were able to adapt their LPD on a trial to trial basis as indicated by the ratio values for the second trials of all conditions (Pooled ratio = 1.12 ± 0.03). However, it is noteworthy that participants overcompensated in the second trial after the weight change. This is especially visible in the SO condition (Ratio 2^nd^ trial SO = 1.21 ± 0.04). In addition, although participants made a predictive error in the first trial after the weight change in the SOLO and SO condition, participants were already able to skilfully lift the object in the second trial after the weight change as can be seen in the significant differences between the first and second trials in the SOLO and SO condition *(all p-values < 0.001)*. This improvement was absent for the EO condition revealing that participants were already able to lift the object accurately in the first trial after the weight change (*p-value = 0.92)*.

Altogether, our results for the light-after-heavy weight changes support the findings of Reichelt et al. that observation of erroneous lifts enables the observer to accurately scale fingertip forces according to the actual object weight (Reichelt et al., 2013). In addition, our results for light-after-heavy weight changes suggest that when individuals observe skilled lifts, they might be able to improve their predictive force scaling although to a lesser extent than after observing erroneous lifts.

### Observers’ lifting force parameters: heavy-after-light weight changes

The right panel of Figure 4 shows the averaged force profiles of the first trial after the object weight changed from light to heavy in a typical subject. When a subject scales his fingertip forces in anticipation of a light object, less force than required (undershoot) will be applied to lift the heavy object adequately (Johansson & Westling, 1988). Figure 4 reveals that the subject upscaled his force generation after observing an erroneous or skilled lift compared to the SOLO condition. In the case of heavy-after-light weight changes, force parameters with ratios smaller than 1 indicate underestimation. As expected, these effects are the opposite for loading phase duration: a ratio value >1 indicates a slower increase in force generation resulting in a longer loading phase duration.

#### Load force rate

*Analysis* of peak load force rate revealed that both main effects of LIFT NUMBER *(F*_*1, 12*_ *= 25.68; p < 0.001)* and CONDITION *(F*_*1, 12*_ *= 5.92; p < 0.01)* as well as the CONDITION X LIFT NUMBER interaction (*F*_*2, 24*_ *= 3.72; p = 0.04)* were significant. Our findings are interpreted in light of the significant interaction effect. As can be seen in Figure 6A, post-hoc analysis revealed that subjects scaled their load forces significantly more accurately in the EO *(Ratio: 0.96 ± 0.05)* and SO conditions (*Ratio: 0.98 ± 0.04*) in comparison with the SOLO condition *(Ratio: 0.82 ± 0.05)* (*p < 0.05)*. These results indicate that observation of both erroneous and skilled lifts allowed participants to anticipatory scale their fingertip forces. When comparing with the second trials after the heavy-after-light weight change, it is noticeable that the first and second trials of the EO condition do not differ significantly *(p = 1)* indicating that participants were already scaling their load forces accurately in the first trial after the weight change. In contrast, in the SOLO and SO conditions, participants significantly upscaled their load forces in the second trial after the weight change (*both p-values < 0.01)*. Importantly, the significant difference between the first and second trials for the SO condition is likely caused by participants overcompensating in the second trial as shown by larger values for this trial (*Ratio: 1.22 ± 0.06)*.

**Figure 6.**
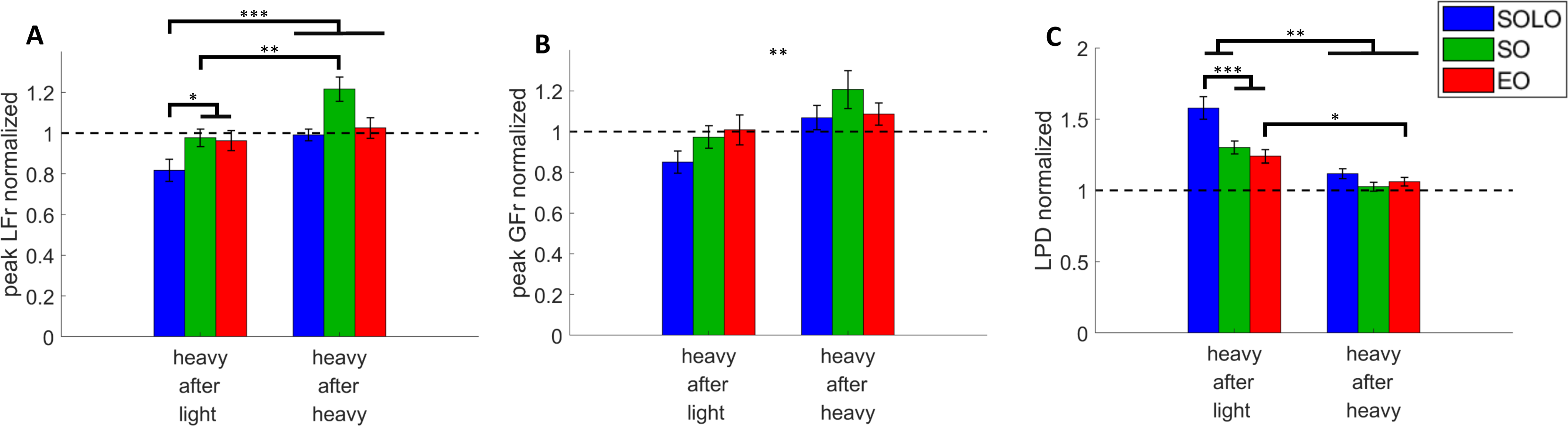
Heavy-after-Light weight changes: Subject group averages for the first and second trials after the weight changed from light-to-heavy for the three conditions [grasping with incorrect weight expectations (SOLO), grasping after observing a skilled lift (SO) and grasping after observing a lifting error (EO)]. **A**. Peak load force rates, **B**. Peak grip force rates and **C**. Loading phase durations. All data is represented as a ratio (normalized to skilled lifting, i.e. heavy-after-light and aeavy-after-heavy divided by the last heavy-after-heavy lift of the same weight sequence block). A ratio > 1 for peak grip force rates and peak load force rates (and a ratio>1 for loading phase durations) indicates that subjects underestimated object weight. All data is presented as the pooled mean ± SEM. *** p > 0.001, ** p > 0.01, * p > 0.05.

#### Grip force rate

A significant main effect of LIFT NUMBER (*F*_*1, 12*_ *= 16.65; p < 0.01)* but not of CONDITION *(F*_*2, 24*_ *= 1.86; p = 0.17)* was found. In addition, the CONDITION X LIFT NUMBER interaction was not significantly *(F*_*2, 24*_ *= 1.44; p = 0.25)*. Accordingly, these results indicate that performance significantly improved after the weight change from the first to the second trial but no differences were found between conditions (Fig. 6B).

#### Loading phase duration

The repeated-measures ANOVA revealed that both main effects and the interaction effect were significant (*all F-values>8.22, all p-values < 0.01)*. As shown in Figure 6C, the loading phase duration in the first trial after the weight change was significantly shortened for both EO and SO conditions (Ratio for EO = 1.24 ± 0.05; Ratio for SO = 1.30 ± 0.04) compared to the SOLO condition (Ratio = 1.58 ± 0.08) *(both p-values < 0.001)*. In addition, the type of action observation (EO vs. SO) did not affect anticipatory force scaling in the first trial after the weight change (*p = 1.00)*. With respect to the second trials after the weight change, it is noticeable that for each condition, participants had a significantly shorter loading phase duration in the second trials after the weight change *(all p-values > 0.05)* indicating that independently of condition, subjects underestimated object weight in the first trial and subsequently improved in the second trial. Lastly, the post-hoc analysis failed to reveal any significant differences between the second trials of the three conditions (*all p-values = 1.00)* indicating that the object internal sensorimotor representation was accurately updated independently of condition.

Altogether, our findings for heavy-after-light weight changes show that observation of skilled lifts enabled participants to improve their predictive force scaling as well as the observation of erroneous lifts.

## Discussion

The present study investigated whether observation of skilled object lifting allows individuals to update their internal sensorimotor representations, which in turn might translate into changes in anticipatory motor control. Importantly, our results not only corroborate recent findings regarding observation of lifting errors (e.g. Buckingham et al., 2014; Reichelt et al., 2013) but also revealed that observation of skilled hand movements can drive predictive motor control, albeit to a smaller extent than observation of explicit movement errors. For this reason, our results not only support the current consensus that grasp observation allows for accurate weight judgement (e.g. Meulenbroek et al., 2007; Shim & Carlton, 1997) but also sheds new light on the role of more implicit, natural, movement cues in mediating motor planning in an observer (Buckingham et al., 2014; Reichelt et al., 2013; Uçar & Wenderoth, 2012).

The first aim of our study was to replicate the results of Reichelt and colleagues. Using a dyadic setting, consisting of a participant and an actor, these researchers revealed that observation of lifting errors can be used to perceive object weight and subsequently allow participants to scale their fingertip forces accurately when lifting the object themselves. When an object with unknown weight was presented, the actor would make a typical lifting error (over- or underestimation of object weight) as he did not have prior knowledge about the object weight (Reichelt et al., 2013). It is plausible that participants deduced object weight based on the observed kinematics: Firstly, it has been well established that over- and underestimation of object weight respectively shortens or elongates the lifting phase when lifting an object (for example see: Gordon et al., 1991; R S Johansson & Westling, 1988). Secondly, Hamilton and colleagues demonstrated that individuals will estimate an object to be light when they observe a short lifting phase and, conversely, will estimate an object to be heavy when observing a longer lifting phase (Hamilton et al., 2007). Our results are consistent with the findings of Reichelt and colleagues: Participants in the current study were capable to predictively scale their fingertip forces with significantly improved accuracy after observing lifting errors indicating a change in object weight. It is noteworthy that in the current study a relatively large overestimation of object weight remained present in the observer after observing the actor lifting the light object erroneously whereas this effect was completely eradicated in the study of Reichelt et al. (Reichelt et al., 2013). This discrepancy might be due to the different movement parameters investigated: While we used force parameters (peak grip and load force rates), Reichelt and colleagues used lifting height as indicator of predictive control. Since peak force rates occur before the time to reach lifting height, it is plausible that subjects can use feedback mechanisms to update their lifting height, but not earlier force parameters, which would therefore be not fully tuned to the current object weight.

The second aim of our study was to test whether observation of skilled lifts mediates predictive motor control as equally as observation of movement errors. With respect to both weight changes, it is interesting to note that predictive scaling of the load force and loading phase after observing skilled lifting significantly improved compared to the SOLO condition but was not as efficient as the observation of lift errors, in particular for light object lifts. This indicates that observation of skilled movement performance can also convey critical information about object weight but to a smaller extent than error observation. It is noteworthy that our results about skilled grasp observation are in contrast with the study of Buckingham et al. Indeed, their study revealed that error, but not skilled lift observation, significantly reduced the learning that is required to grasp a novel, surprisingly light object (Buckingham et al., 2014). Importantly, there are two major considerations to take into account while comparing the results of the Buckingham study and ours. Firstly, while we used two differently weighted object with identical appearance, Buckingham and colleagues used two objects that were identical in weight but different in size (i.e. ‘Size-Weight Illusion’). It is likely that this size difference caused a strong initial bias regarding weight expectations towards the objects (for example see: Gordon et al., 1991; Peters, Ma, & Shams, 2016). Secondly, in the Buckingham study, participants were not familiarized with the objects prior to observing object lifting videos. This lack of familiarization and the presence of a size-weight illusion might cause a different action observation strategy for extracting information from skilled or erroneous lifting: When lifting skilfully, the kinematics of the lifting phase tend to have a similar duration regardless of object weight (Gordon et al., 1991). According to this, it is likely that participants presumed that weight and size were congruent when observing skilled object lifting, therefore leading them to focus mostly on the size cue. In contrast, the observation of lifting errors reveals an incongruence between size and expected weight which likely led participants to not only focus on size but also on the movement kinematics. In our study, participants could only rely on the observed movement kinematics to assess object weight as we excluded other visuals cues indicating object weight. Interestingly, participants could estimate object weight during both the observation of skilled and erroneous lifting. For observation of errors, it is likely that participants perceived object weight by focusing on the lifting phase duration and grasp duration (hand-object contact without movement) (Hamilton et al., 2007; Roland S. Johansson & Flanagan, 2009). Having experienced these typical lifting errors in the SOLO condition, participants were likely to interpret the lifting errors made by the actor and adjust their internal sensorimotor representation accordingly. In contrast, when observing skilled lifting, the kinematic profiles of a heavy or light lift are more similar compared with lifting errors on the same objects. It is therefore possible that participants developed an observational strategy emphasising on other parameters to differentiate between weights such as the hand contraction state (Alaerts, Senot, et al., 2010; Uçar & Wenderoth, 2012), the reaching phase (Ansuini, Cavallo, Bertone, & Becchio, 2014) or the intention of the actor (Cavallo, Koul, Ansuini, Capozzi, & Becchio, 2016; Wasmuth & Lima, 2016).

The neural substrate responsible for the sensorimotor mapping of observed actions into one’s own motor repertoire is likely to be supported by the ‘mirror neuron system’, located in a subset of sensorimotor brain areas (Giacomo Rizzolatti & Craighero, 2004). Mirror neurons were found to discharge both when a monkey performs a goal-directed hand action and when observing another individual performing the same action (Buccino, Binkofski, & Riggio, 2004; di Pellegrino, Fadiga, Fogassi, Gallese, & Rizzolatti, 1992; G Rizzolatti, Fadiga, Gallese, & Fogassi, 1996; Umiltà et al., 2001). More importantly, it has recently been demonstrated that the action observation-induced increase of excitability in the primary motor cortex, so called ‘motor resonance’, reflects specific parameters during grasp observation such as the hand contraction state (Alaerts, Swinnen, & Wenderoth, 2010) or observed movement kinematics, indicating object weight (Alaerts, Swinnen, et al., 2010; Senot et al., 2011), object shape (Buckingham et al., 2014) and even the intentions of the observed actor (Wasmuth & Lima, 2016). In the current study, participants were not able to perceive object weight via intrinsic object properties. For this reason, it is plausible that participants had access to information about object weight by mapping onto their own motor repertoire observed visuomotor cues such as object kinematics and hand contraction states.

In conclusion, participants in the present study were familiarized to two different object weights and generated a sensorimotor repertoire for skilled lifting (by applying accurate forces following consecutive lifts of a same object) and for lifting errors (by over- or underestimating forces after a weight change). After this initial process, participants lifted objects in turns with an actor. In this dyadic setting, the only way individuals could extract information about weight, and in turn plan their subsequent motor command, was by embodying the observed visuomotor cues into their own sensorimotor repertoire. Our results not only support recent findings regarding the effect of observation of explicit movement errors on mediating predictive motor control but also highlight that the observation of skilled movements, carrying more implicit visuomotor cues, can also drive motor planning. Interestingly, anticipatory force scaling in the first trial following skilled lift observation was not as accurate as following error observation, and still improved in the second trial. This highlights that different action observation mechanisms could contribute to mediating anticipatory motor control in an observer when surprising or erroneous movements are performed (Cretu, Ruddy, Germann, & Wenderoth, 2019).

## Acknowledgements

This work was funded by the Research Foundation Flanders (FWO) Odysseus Project (Fonds Wetenschappelijk Onderzoek, Belgium) awarded to M.D. The authors declare no competing financial interests.

